# Lipid type doping of the sponge (L_3_) mesophase

**DOI:** 10.1101/2021.02.22.432284

**Authors:** Christopher Brasnett, Adam Squires, Andrew Smith, Annela Seddon

**Affiliations:** School of Physics, University of Bristol, Tyndall Avenue, BS8 1FD, UK; Bristol Centre for Functional Nanomaterials, School of Physics, University of Bristol, Tyndall Avenue, BS8 1FD; Department of Chemistry, University of Bath, Bath, BA2 7AY, UK; Diamond House, Diamond Light Source Ltd., Harwell Science and Innovation Campus, Fermi Ave., Didcot OX11 0DE, U.K

**Author notes:** Electronic Supplementary Information (ESI) available: [details of any supplementary information available should be included here]. See DOI: 10.1039/cXsm00000x/.

## Abstract

The polymorphism of lipid aggregates has long attracted detailed study due to the myriad factors that determine the final mesophase observed. This study is driven by the need to understand mesophase behaviour for a number of applications, such as drug delivery and membrane protein crystallography. In the case of the latter, the role of the so-called ‘sponge’ (L_3_) mesophase has been often noted, but not extensively studied by itself. The L_3_ mesophase can be formed in monoolein/water systems on the addition of butanediol to water, which partitions the headgroup region of the membrane, and decreases its elastic moduli. Like cubic mesophases, it is bicontinuous, but unlike them, has no long-range translational symmetry. In our present study, we show that the formation of the L_3_ phase can delicately depend on the addition of dopant lipids to the mesophase. While electrostatically neutral molecules similar in shape to monoolein (DOPE, cholesterol) have little effect on the general mesophase behaviour, others (DOPC, DDM) significantly reduce the region in which it can form. Additionally, we show that by combining cholesterol with the anionic lipid DOPG, it is possible to form the largest stable L_3_ mesophases observed to date, with correlation lengths over 220 Å.

## 1 Introduction

One of the principle motivations for the study of self-assembled lipid systems is their astonishing range of potential applications, ranging from templating and drug delivery, to membrane protein crystallisation^1–6^. Of these, membrane protein crystallisation using the so-called *in meso* or lipid cubic phase (LCP) method has long been cited as a principal source of motivation for studies of lipid polymorphism^7–15^.

Self-assembled lipid aggregates can exhibit a number of different symmetries shown in Figure 1a)-d), from the planar 1 dimensional lamellar bilayer stacks (L_α_), 2 dimensional arrays of hexagonally arranged cylinders, or 3 dimensional cubic phases. In cubic phases, the lipid bilayer spans a triply periodic minimal surface, a surface defined by having zero mean curvature at all points, and separates the system bicontinuously into two water channels^16^. Three bicontinuous cubic phases are known, the Primitive 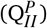, Diamond 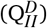, and Gyroid 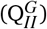, in order of increasingly negative Gaussian curvature. In addition to these extensively-studied mesophases, the sponge (L_3_) mesophase is occasionally observed upon the addition of other molecules to the monoolein/water system. Mesophases have characteristic small angle X-Ray scattering (SAXS) patterns according to their topology and size^17^. In the case of the L_3_ mesophase, the characteristic broad peak measured using SAXS measures the interbilayer correlation length.

**Fig. 1.**
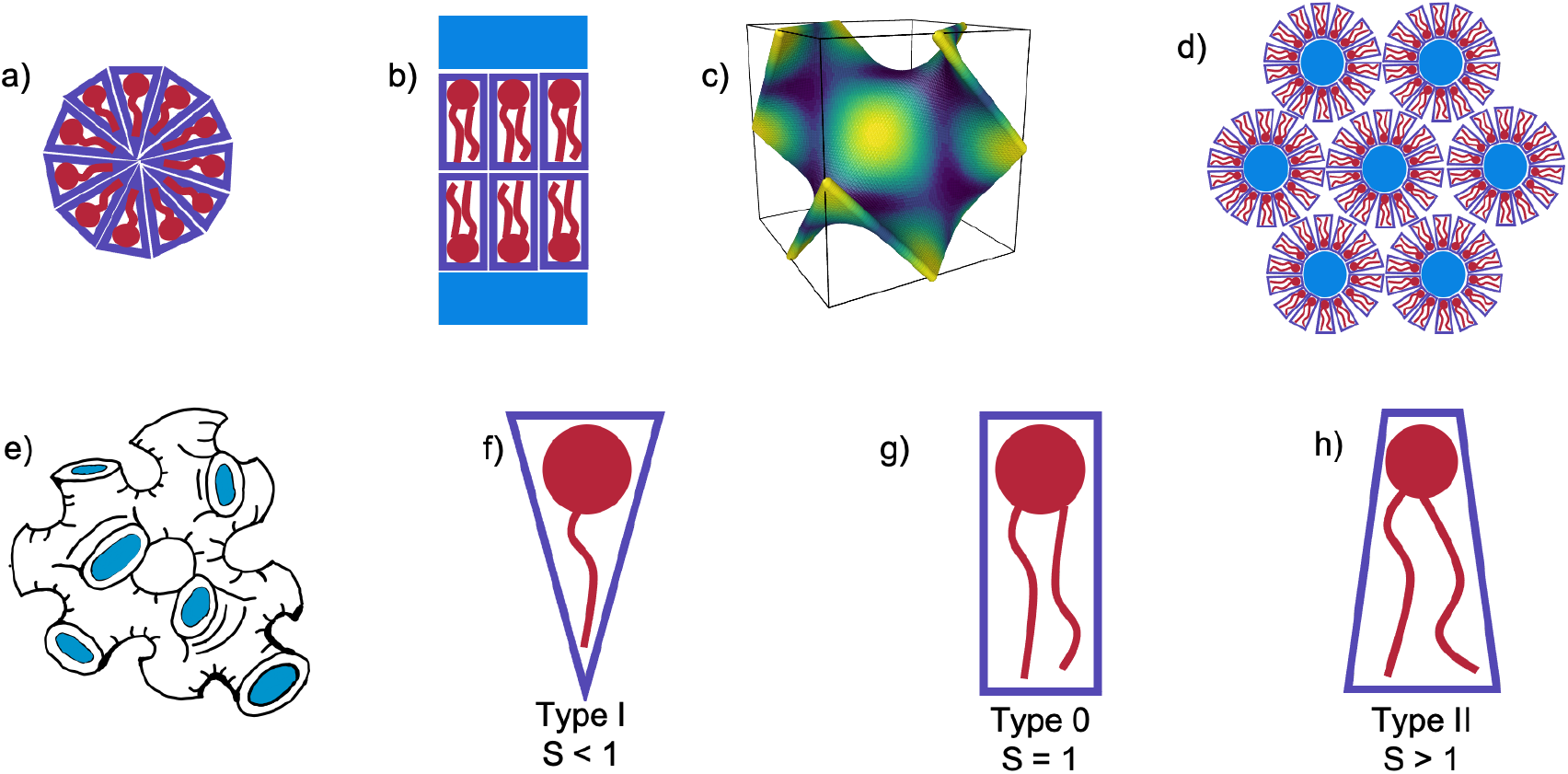
Illustrations of possible self-assembled mesophases and packing types of lipids and surfactants. a) a micelle, b) a flat bilayered L_α_ mesophase surrounded by water channels, c) a 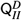 minimal surface (coloured according to Gaussian curvature: light patches flat and dark more curved), d) a Hexagonal (H_*II*_) mesophase, with blue water channels extending in and out of the page, e) a sponge (L_3_) mesophase with water channels highlighted in blue, f) A type I molecule, with S < 1, g) a type 0 molecule with S = 1, h) a type II molecule with S > 1.

The sponge phase is known to be closely related to the cubic phase, similarly consisting of a lipid bilayer separating bicontinuous water channels, but without long-range translational symmetry^18–21^. Cherezov *et al.* showed that a number of additives to the monoolein (MO)/water system can form the sponge phase in lipid systems, a common feature among one class in particular being that they are small amphiphiles with a number of both hydrogen bond acceptors and donors^22^. This enables them to interact with both the water channels of the 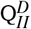 mesophase as well as the hydrophobic region of the membrane, later confirmed in ^1^H NMR studies by Evenbratt *et al.* ^23^. The net effect of these interactions is the swelling of the 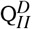 mesophase as the interface is partitioned by the additives and flattened. Beyond a certain concentration when the bending moduli of the membrane is lowered, the 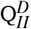 mesophase becomes disordered and the system becomes a L_3_ mesophase. Upon further increase of the sponge-forming additive, a flat L_α_ mesophase will emerge.

One of the greatest successes of the LCP technique to date has been the solution of the structure of the human *β*_2_ adrenergic G-protein-coupled receptor^24,25^. The high-throughput screening techniques used in these studies for finding successful crystallisation conditions noted that including a significant proportion of cholesterol in the mesophase was essential for crystal growth. In the case of crystallisation of the entire protein complex, crystals were harvested from a ‘sponge-like mesophase’. While the mesophase was not explicitly characterised, the addition of PEG400 is known to induce this transition, so it is likely that this was the case^26–28^. Similarly, crystallisation of human microsomal prostaglandin E2 synthase 1 by Li *et al.* found that doping the monoolein membrane at a level of 5% mol with DOPC, a zwitterionic phospholipid, was essential to guarantee successful crystallisation ^29^.

To add to the significance of the sponge phase, other membrane proteins had previously been crystallised from the sponge phase directly, but the solution of the human *β*_2_ adrenergic G-protein-coupled receptor demonstrates the significance of the method^30–32^. Understanding the exact mechanisms of LCP remains an area of extensive study, with Zabara *et al.* demonstrating that the sponge phase can play an important intermediary role in the crystallogenesis of membrane protein crystals^33^.

The mesophase behaviour of monoolein has been extensively studied on its own, forming a 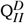 mesophase in excess water at room temperature, and undergoing a transition to the H_*II*_ mesophase when heated^34,35^. In addition to studying the mesophase behaviour of monoolein alone, many studies have additionally investigated the effect of lipid type doping on the structure of the self-assembled behaviour of lipid systems, and for an excellent review we refer the interested reader to the work of van ‘t Hag *et al.* ^14^. Lipid type refers to the categorisation of lipids and surfactants according to their packing parameter, *S*^36^:

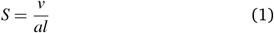

where *v* is the volume of the hydrocarbon tails, *a* the interfacial surface area, and *l* the maximum effective tail length. Molecules can then be catagorised depending on their value of S, as we show in Figure 1f)-h), which is a key determinant in the self-assembled mesophase that results from aggregates of molecules of that type. For S < 1, the interfacial area dominates, and a positively curved membrane where the membrane curves away from its hydrophobic region results. Conversely, molecules with S > 1 will have negatively curved interfaces. For S = 1, cylindrical-like molecules will become result in planar bilayered systems. When added to cubic phases, Cherezov *et al.* showed that generally, the cubic phase can hold up to around 20% mol doping of an additional lipid before reverting to the preferred mesophase of the dopant^37^.

One of the main barriers to the success of the LCP method is the small water channels of the cubic phase ^38^. Attempts to overcome this have often used lipids with charged headgroups in order to promote intra-bilayer electrostatic repulsion as to increase the size of the lattice parameter, and flatten the cubic phase observed to a 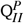 ^12^. However, our recent work has demonstrated that the addition of common salts at low concentrations will screen intra-bilayer charge repulsion, and so revert the mesophase of the lipid system back to the 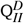 ^39^.

Noting that the adaptability of the sponge phase is a little-understood area, in the present work, we investigate lipid type doping of the sponge phase, using several common lipid and detergent additives. The possibility of doping the sponge phase has been noted in direct sponge phase crystallisation trials, where the use of 1% w/w cholesterol was necessary for successful crystallisation of a bacterial photosynthetic core complex^31^. While it would be ideal to screen every possible crystallisation condition for the mesophase behaviour, we can seek to understand a broader set of design rules by understanding the conditions under which the L_3_ mesophase forms. We investigated the effect on the 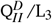 transition on doping monoolein with several common additive molecules of different packing parameters: dioleoylphosphoethanolamine (DOPE), cholesterol, dioleoylphosphocholine (DOPC), and n-Dodecyl *β*-D-maltoside (DDM). In addition, we looked at using a combination of the anionic dioleoylphosphoglycerol (DOPG) and cholesterol together, after Tyler *et al.*, to maximise the lattice parameters obtained^12^.

## 2 Materials and Methods

Monoolein (MO) was received as gift from Danisco and used without further preparation. Cholesterol and DDM were purchased from Sigma Aldrich in powdered form. 1,4-butanediol was purchased from Sigma Aldrich. DOPC, DOPE, DOPG were purchased from Avanti Polar Lipids in powdered form. Lipids (aside from DDM) were prepared in dichloromethane at concentrations of 0.1 M, and mixed at the required doped molarities. DDM was prepared in ethanol at a concentration of 0.05 M. Monoolein was prepared at both 0.1 M and 0.05 M concentrations, and 2.5%, 5%, 7.5%, and 10% mol doped monoolein solutions were then prepared volumetrically. 70 *μ*l of doped lipid mixtures in solvent were transferred to 1.5mm X-Ray capillaries (Capillary Tube Supplies UK Ltd.), and left to evaporate for 3 days. The remaining solvent was removed under vacuum, leaving a film of dried mixed lipid on the capillary walls, before the addition of 50 *μ*l of lyotrope. The capillaries were then sealed, and put through 3 freeze-thaw cycles to ensure the sample was at equilibrium before measurement. Preparation of x-ray capillaries under vacuum is known to have significant effects on mesophase behaviour, so this last step ensured that no out of equilibrium effects were measured^40^.

We chose to measure the self-assembled mesophase at 9 volume/volume proportions of butanediol/water lyotrope content: 0%, 20%, 32.5%, 35%, 37.5%, 40%, 42.5%, 45%, 47.5%. For monoolein alone, the 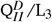 transition is known to occur at 30% v/v butanediol, and the L_3_/L_*α*_transition at 50% v/v^22^. This selection of lyotropes therefore allows us to understand the mesophase behaviour in i) water alone, ii) a point significantly below the expected transition, in case any significant anomalous behaviour is observed, and iii) a granular range of lyotropes in a region where the sponge phase is known to exist, in order to observe the lowering of the second transition.

SAXS measurements were performed on a SAXSLAB Ganesha 300XL instrument with a q range of 0.015–0.65 Å^-1^ for 600s per sample. All scattering patterns measured are plotted in the SI. Mesophases and their sizes were determined from their characteristic Bragg peak spacings measured with SAXS, as detailed in S1 of the SI. Bragg peaks were found using Python scripts written in-house and available at https://github.com/csbrasnett/lipidsaxs.

## 3 Results and Discussion

### 3.1 Monoolein sponge phases

In Figure 2, we validate the well-known result of Cherezov *et al.* of the mesophase behaviour of the MO/water/butanediol system^22^. On the initial addition of butanediol to a proportion of 20% v/v of the lyotrope, the size of the 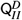 mesophase swells from 97 Å to 123 Å in water. In our focused range between 32.5% and 47.5% butanediol, we observe a sponge phase with a size increasing linearly from 100 Å to 154 Å.

**Fig. 2.**
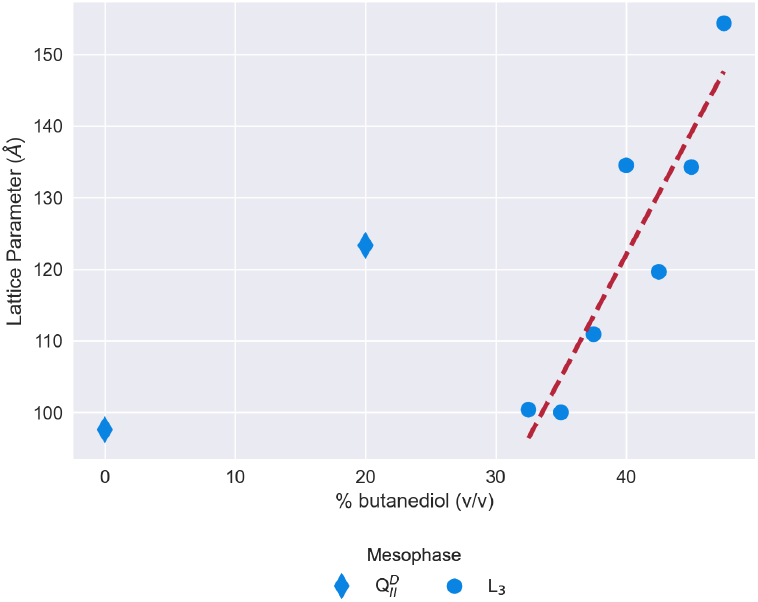
The mesophase and size behaviour of a system of monoolein and a lyotrope of butanediol and water, with increasing proportions of butanediol in the lyotrope. The shape of the scatter point indicates the mesophase, a Diamond for the 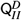 mesophase and a circle for the L_3_ mesophase. The red dashed line is a linear regression to the L_3_ mesophase data as a guide

To form the L_3_ meosphase, Cherezov *et al.* use a set ratio of lipid:lyotrope of 60:40 % weight/weight, the excess water point of monoolein in water^22,35^. To validate our excess lyotrope method, we measured the correlation lengths of sponge phases at 3 excess weight ratios, and observed no significant change (±10 Å, see Figure S1 in SI). This demonstrates that the results throughout this work will be valid for any excess hydration conditions.

### 3.2 Effect of single dopants on the formation of the L_3_ mesophase

#### 3.2.1 Type II: DOPE and cholesterol

Figure 3 shows how the doping of monoolein with Type II lipids affects the phase behaviour and size of the self-assembled mesophase as the proportion of butanediol in the lyotrope is increased. For a simple ternary MO/H_2_O/butanediol system, the 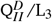 phase change is expected at 30% v/v butanediol. In both the DOPE- and cholesterol-doped systems, the 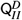 phase swells from a lattice parameter of around 100 Å to between 110 Å and 120 Å on an initial increase of butanediol in the lyotrope to 20% v/v. We then increase the proportion of butanediol in the system to 32.5% v/v, at which point we see a difference between the two dopants. While the cholesterol-doped systems continue to swell the 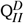 phase to a size of around 130 Å, the DOPE-doped systems have undergone a transition to the L_3_ phase, similar to pure monoolein. Cholesterol is known to condense and stiffen membranes, so while it may also be a type II molecule, the stiffer membrane does not favour undergoing a phase transition at this point^41–44^.

**Fig. 3.**
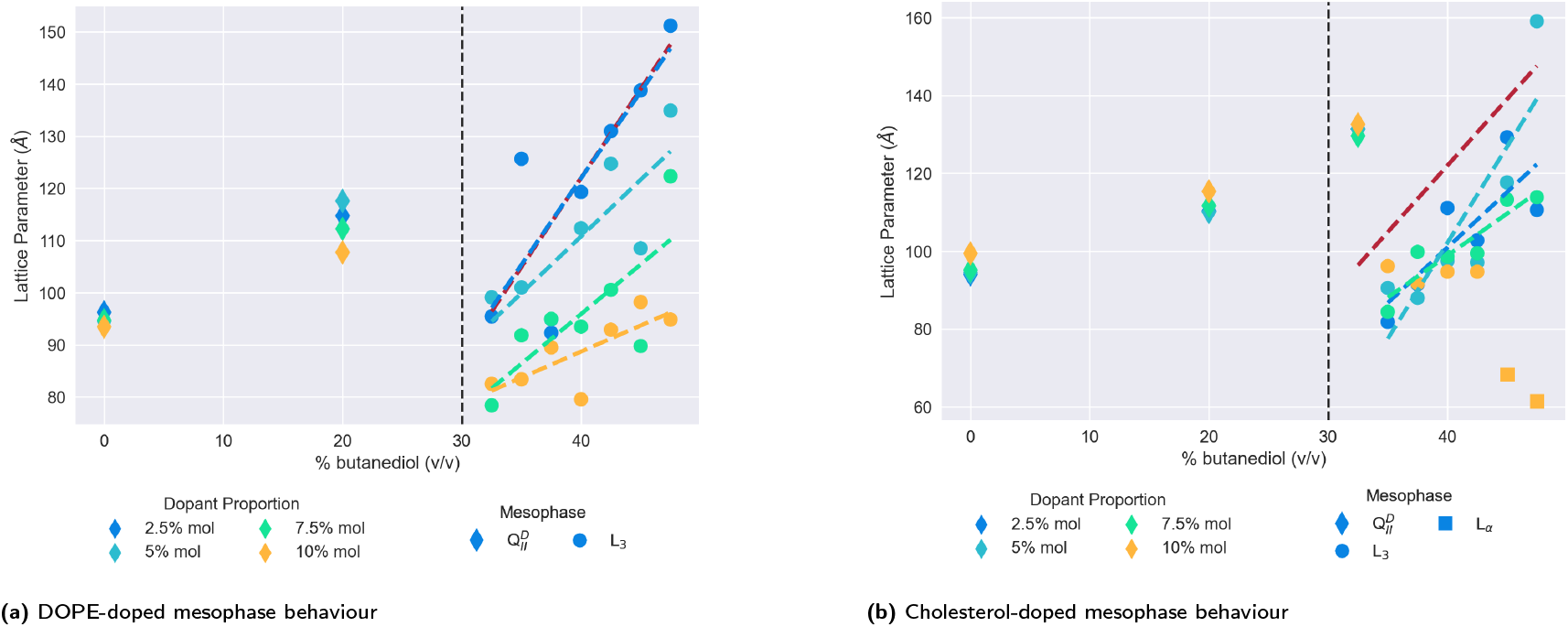
The mesophase and size behaviour of type II doped monoolein systems changing as butanediol is introduced into the system. Molar proportion of dopant is indicated by colour: 2.5% (Blue), 5% (Turquoise), 7.5% (Green), and 10% (Yellow). The shape of the scatter point indicates the mesophase: 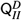 (Diamond), L_3_ (Circle), L_α_ (Square). Dashed lines have been fitted using linear regression as a guide for the trend of the correlation length of the L_3_ mesophases, and coloured accordingly. In addition, the Red dashed line is the linear regression to the pure monoolein L_3_ mesophases seen in Figure 2 as a reference. The vertical black dashed line at 30% v/v indicates the position of the 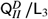 transition in a pure monoolein lipid system.

A further increase in the butanediol content of the lyotrope to 35% results in both the DOPE- and cholesterol-doped systems transitioning to L_3_ mesophases. In both cases, the transition sees the correlation length of the L_3_ mesophase reduced in comparison to monoolein alone. There is no clear trend as to how the increase in dopant proportion immediately changes the size of the L_3_ correlation length, in both the DOPE and cholesterol. However, the subsequent trend shows that in DOPE, the size of the L_3_ phase increases more when the quantity of dopant is lower. Indeed, at 47.5% v/v butanediol, there is a decrease from 151 Å to 94 Å of the L_3_ correlation length on increasing the dopant proportion from 2.5% to 10% mol.

Unlike DOPE, the cholesterol-doped L_3_ phases do not demonstrate a clear trend between L_3_ correlation lengths and dopant proportions. Additionally, most of the sponge phases (aside from an anomalous 5% mol doped at 47.5% v/v butanediol) are smaller than DOPE-doped ones. This could similarly be explained by the stiffening effect that cholesterol has on the membrane. A recent study by Chakraborty *et al.* showed that as the cholesterol content of vesicles is increased, both the membrane thickness and relative bending rigidity moduli do so correspondingly^41^. In the context of this work, a thickening effect on the membrane would explain why the L_3_ correlation length is mostly reduced in comparison to the L_3_ phases we see doped with DOPE.

The principal difference between the systems doped with either DOPE or cholesterol is the emergence of the L_α_ mesophase at high concentrations of cholesterol. As shown in Figure 3b, at a level of 10% mol cholesterol doping, the system undergoes a second phase transition from the L_3_ to the *L_α_* at a butanediol lyotrope concentration of 45% v/v, whereas no such transition happens in systems doped with DOPE. Cherezov *et al.* found that above a 20% mol doping, cholesterol-doped monoolein will form a flatter 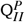 mesophase^37^. In contrast, doping with DOPE above a 20% mol level will results in a more curved H_*II*_ mesophase. Although at 10% mol doping in water, both DOPE and cholesterol doping should be stable in the 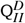 mesophase, the tendency towards becoming more curved and flatter respectively are reflected in the lowering of the L_3_/L_α_ transition at 10% mol cholesterol.

#### 3.2.2 Type 0: DOPC

In contrast to the relatively simple and unmodified phase sequences that are observed by doping with type II lipids, the mesophase behaviour for monoolein doped with the type 0 lipid DOPC is significantly more complex. We show the results in Figure 4, split between lower levels of doping (Figure 4a) and higher levels (Figure 4b). As one expects, the initial introduction of butanediol into the lyotrope swells the size of the 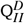 mesophase observed.

**Fig. 4.**
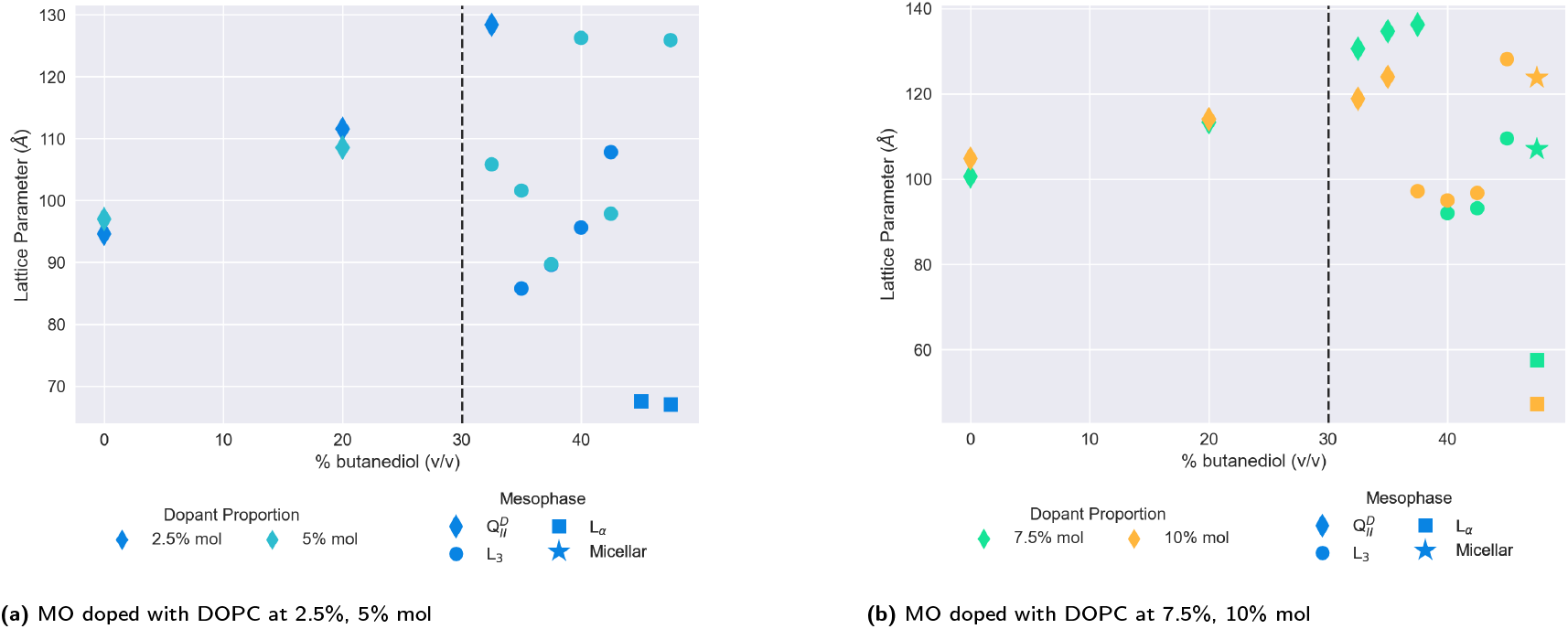
The mesophase and size behaviour of MO with a) low doped proportions (2.5%, 5% mol) and b) high doped proportions of DOPC (7.5%, 10% mol) with varying proportions of butanediol in the lyotrope. The molar proportions are indicated by the colour of the scatter points, and the mesophase by their shapes. The dashed vertical black lines indicate the location of the 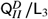 transition in a system of MO alone.

At lower proportions of DOPC in the membrane(Figure 4a), the 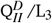 transition does not appear to shift significantly with respect to the point for monoolein L_3_ phases alone. However, the range in which the phase exists is narrowed: at a 2.5% mol doping, the L_3_/L_α_ transition is reduced to 45% v/v butanediol from 50% v/v. At DOPC has a larger headgroup as a type 0 lipid, and forms L_α_ mesophases of its own accord, this is to be expected. Interestingly, the same transition is not observed in this range for the 5% mol doped systems, which was observed to be a L_3_ mesophase up to 47.5% v/v butanediol in the lyotrope. For all the L_3_ mesophases observed in the lower doped DOPC systems, the observed correlation lengths are smaller than for either monoolein alone, or for the L_3_ mesophases seen in the type II systems.

The results for higher dopant proportions of DOPC are plotted in Fig 4b. On increasing the proportion, the 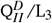 transition point is significantly increased to 37.5% v/v butanediol for 10% mol DOPC, and 40% v/v butanediol for 7.5% mol DOPC. Molecular dynamics and spectroscopy studies investigating the water/lipid interface have shown that phosphocholine headgroups allow for a looser packing of water at the interface, and that hydrogen bonds can be formed between the phosphocholine carbonyl groups and water^45–49^. Evenbratt *et al.* showed using ^1^H NMR that diols form the L_3_ phase by molecular partitioning at the polar/non-polar interface^23^. The introduction of high levels of DOPC therefore increases the hydration of the bilayer at the interface, at the expense of the presence of butanediol, and so the 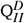 mesophase is sustained for higher proportions of butanediol in the lyotrope than is otherwise expected.

Considering the increased hydration of the interface with the introduction of more DOPC, it is perhaps surprising that the L_3_/L_α_ transition is not reduced correspondingly. We only observed a system of coexisting micelles and L_α_ mesophases for both dopant levels at 47.5% v/v butanediol. As Figure 4b shows, for both the higher proportions of DOPC, we only observed a L_3_/L_α_ transition at 47.5% v/v butanediol for both systems, where there is also coexistence with a micelles. As a type 0 lipid, DOPC has a larger headgroup to begin with than monoolein, which could be expected to lower this transition accordingly. That the transition is only lowered slightly suggests that there is a delicate interplay in this transition between the role of headgroup size and the partitioning effect of butanediol.

#### 3.2.3 Type I: DDM

The addition of DDM to monoolein in water at sufficient concentration is known to destabilise the 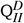 mesophase induce a transition to the *L_α_* phase^50–52^. A 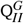 mesophase is observed at intermediate concentrations, as a result of molecular shape mismatch: the 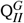 mesophase has the largest interfacial surface area of any of the cubic phases^17^.

We plot the results for lower- and higher-doped systems in Figure 5a and 5b respectively. In contrast to DOPE, cholesterol, or DOPC, the addition of butanediol to the lyotrope in lower-doped systems does not have any significant swelling effect on the 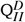 lattice parameter, although the addition of 5% DDM itself increases the size of the 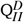 seen to 145 Å, which is maintained at 20% v/v butanediol. Above 30% v/v butanediol, we observed a transition to the L_3_ mesophase for both 2.5% mol and 5% mol doped systems. As the proportion of butanediol is further increased, the mesophase behaviour of the lower-doped systems diverges. The 2.5% mol doped system continues in the L_3_ phase, while for the 5% mol doped system, the L_3_/L_α_ mesophase transition is significantly lowered to 35% v/v butanediol, where it coexists with micelles. In comparison, the 2.5% mol system only sees this transition at 47.5% v/v butanediol, the same lyotrope conditions for very highly-doped systems of DOPC (Figure 4b).

**Fig. 5.**
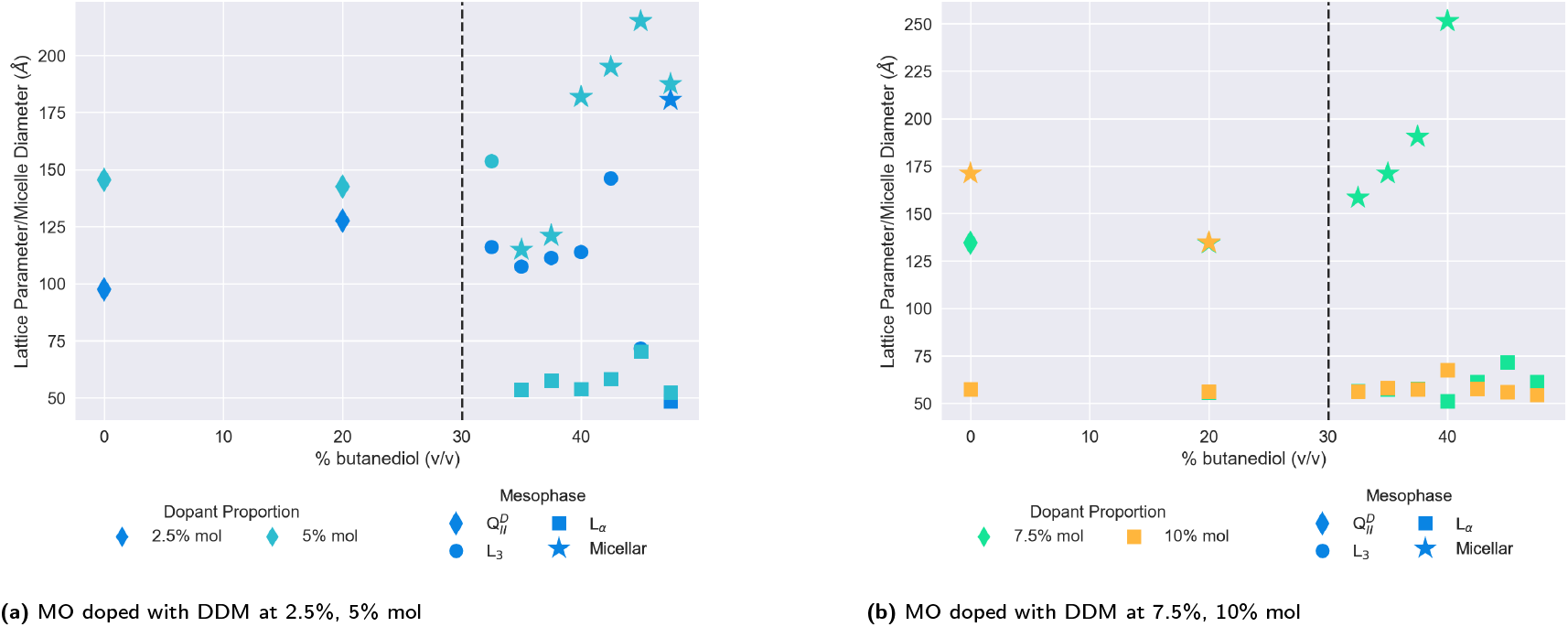
The mesophase and size behaviour of MO doped with a) low (2.5%, 5% mol) proportions of DDM, b) high (7.5%, 10% mol) proportions of DDM with varying proportions of butanediol in the lyotrope. The dopant proportion is indicated by the colour of the scatter point, and the mesophase by its shape. The vertical dashed lines indicate the location of the 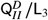 transition in a system of only MO.

On further increasing the proportion of DDM in the membrane more, the destabilising effect of DDM is observed immediately. While at 7.5% mol, the system in water remains in the 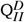 mesophase, at 10% mol, the system has transformed into a coexisting L_α_ and micellar system. The 7.5% mol system also exhibits this at 20% v/v butanediol, and this is subsequently observed for both strongly-doped systems above 30% v/v.

### 3.3 Using electrostatic forces to form sponge phases

In addition to the 4 dopants described above, we screened the anionic lipid DOPG as a possible dopant to form the L_3_ mesophases. As a dopant in MO/water systems, DOPG forms the 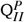 mesophase by introducing intra-membrane repulsion^12^. The 1D SAXS profiles of these experiments are in Figures S19-S21 of the SI show no discernible mesophase structure, indicating that the presence of electrostatic forces alone can prohibit self-assembly into a distinct mesophase.

In order to overcome the destruction wrought by the intramembrane electrostatic repulsion from DOPG alone, we doped a monoolein mesophase using both DOPG, and cholesterol. A tertiary system of MO, DOPG, and cholesterol was previously used by Tyler *et al.* to create some of the largest lipid cubic phases observed, showing that a mixture of 80/15/5 MO/cholesterol/PG heated to 45°C adopts a 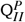 phase with a lattice parameter of 415 Å. Considering our work showing that the L_3_ phase is unstable with even very low quantities of electrostatic doping - and is remarkably stable with even significant levels of cholesterol - we chose to use two high ratios of cholesterol to DOPG while keeping the proportion of MO the same in order to maximise chances of observing the L_3_ mesophase in the lyotrope sequence.

We show both sets of results in Figure 6. A L_3_ mesophase was only observed in the system with 1% mol DOPG and 9% mol cholesterol. On changing this ratio to 3% mol and 7% mol respectively, we observed extremely large L_α_ mesophases, in one instance with a lattice parameter of over 500 Å, and consistently over 200 Å. Where we did observe the L_3_ phase in the first system, it had a consistently larger correlation length than we observe for MO alone in Figure 2. Over the lyotrope range from 32.5% v/v butanediol to 47.5%, the correlation length grows from 110 Å to 222 Å, with a peak of 239 Å observed at 42.5%. Therefore, the condensing effect of cholesterol previously observed in Figure 3b can be used advantageously to stabilise electrostatically-doped systems to result in the largest L_3_ mesophases observed to date.

**Fig. 6.**
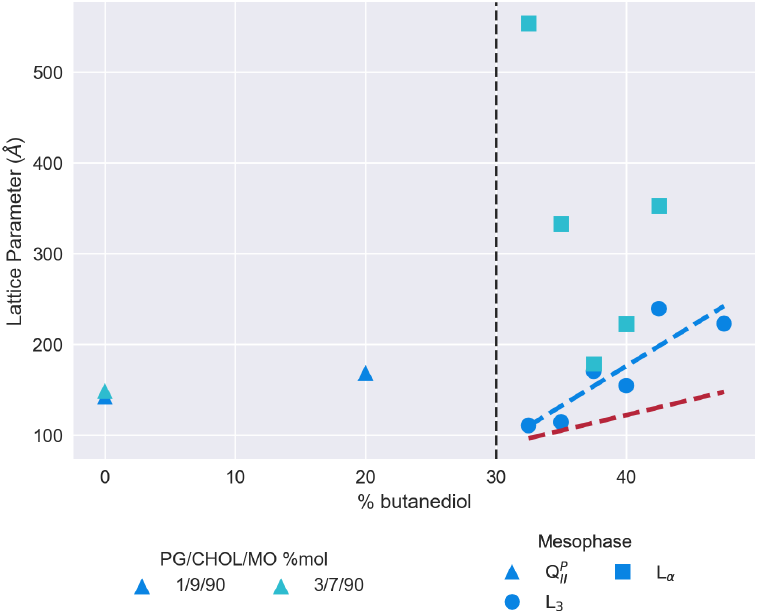
Mesophase and size behaviour of monoolein doped with DOPG and cholesterol with a lyotrope of increasing butanediol content. The dashed red line shows the trend for the MO/water/butanediol system measured in figure 2, and the dashed blue line a linear regression to the L_3_ mesophase data for the 1/9/90 DOPG/cholesterol/MO % mol system. The vertical dashed black lines indicate the location of the 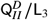 transition for an undoped MO lipid system.

## 4 Conclusions

We have demonstrated that the lipid L_3_ sponge mesophase can successfully and stably be doped with additive lipids. In most cases, we have shown that the presence of additive lipids shrinks the correlation length of the L_3_ mesophase. This finding will have significance for designing LCP trials, as proteins with both large extracellular domains and essential co-crystallisation lipids will need to have the membrane environment carefully tailored to maximise the chances of successful crystallisation. However, we have also shown that use of electrostatic lipids such as DOPG can be used in conjunction with cholesterol to significantly increase the size of L_3_ mesophases. Using these cholesterol-stabilised electrostatically-doped mesophases, we have observed sponge phases with correlation lengths of around 240 Å, significantly larger than in MO alone. These results will should help inform the engineering design rules for future LCP crystallisation trials.

Regarding the 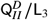 transition itself, here we have shown that the transition is sensitive to both the average headgroup area of the membrane, and the bending modulus. In the case of the former, increasing the interfacial area per molecule in the system increases the proportion of sponge-forming agent required in comparison to L_3_ mesophases formed of MO alone. Using cholesterol - known to increase membrane moduli - slightly increases the proportion of sponge-forming agent required. DDM, which destabilises lipid cubic phases, drastically reduces the ability for L_3_ mesophases to form. Techniques used to study the L_3_ mesophase in recent years have mainly been limited to SAXS, such that the transition and relationships between the 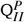 and the L_3_ mesophases are studied statically. In future, methods to study the L_3_ mesophase could take into account the dynamic nature of the phase transition, by investigating the structural rearrangement of lipid molecules in the process through techniques such as molecular dynamics.

In the context of LCP cyrystallisation, this study has shown that while many previous studies have been able to dramatically increase the lattice parameter of cubic phases using anionic doping, the effect of other crystallisation additives can prove to be otherwise counter productive. While it is impractical to investigate the combined effect of every possible crystallisation screen, we have shown here that in particular, the inclusion of cholesterol can stabilise electrostatically charged systems. These results should help inform future LCP trials where the need for large water channels is key to successful crystallisation conditions.

## SI

Supplementary information contains all scattering patterns for the data discussed, and details on structural parameter calculations.

## Acknowledgements

The Ganesha X-ray scattering apparatus used for this research was purchased under EPSRC Grant ‘Atoms to Applications’ Grant ref. EP/K035746/1.

CB acknowledges funding from EPSRC EP/N509619/1.

We are grateful to Diamond Light Source for beam time awards (SM17767-1) to this project and to the staff, particularly Nick Terrill, on beamline I22 for their support.

