## Supplemental Information for "Lipid type doping of the sponge (L_3_) mesophase"

#### S1 SAXS measurements

Lipid mesophases have characteristic scattering patterns, with Bragg peaks appearing in the following ratios:

- $x_{(n,L_\alpha)}$: $1 : 2 : 3 : \ldots$
- $x_{(n,H_{II})}$: $1 : \sqrt{3} : \sqrt{4} : \sqrt{7} : \ldots$
- $x_{(n,Q^D_{II})}$: $\sqrt{2} : \sqrt{3} : \sqrt{4} : \sqrt{6} : \sqrt{8} : \sqrt{9} : \ldots$

so that the lattice parameter can be calculated as:

$$a_{(n,L_\alpha)} = \frac{2\pi}{q_n} \times x_{(n,L_\alpha)}$$
$$a_{(n,H_{II})} = \frac{2}{\sqrt{3}}\frac{2\pi}{q_n} \times x_{(n,H_{II})}$$
$$a_{(n,Q^D_{II})} = \frac{2\pi}{q_n} \times x_{(n,Q^D_{II})}$$

The sponge phase has a broad peak, the centre ($q_c$) of which defines a bilayer-bilayer correlation length:

$$a = \frac{2\pi}{q_c} \tag{1}$$

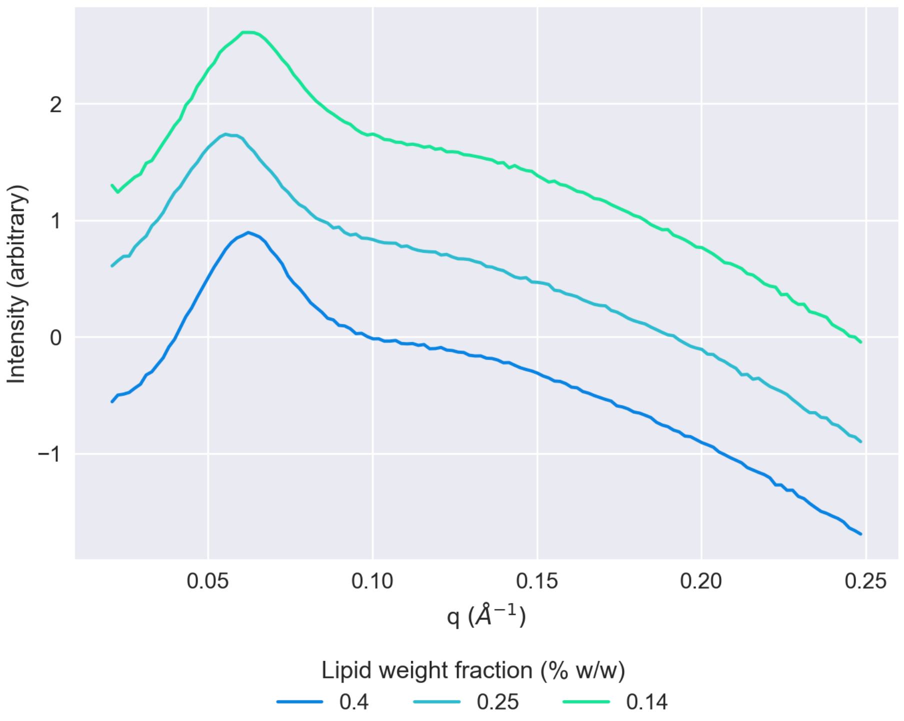


Figure S1: 1D SAXS patterns showing different weight ratios of lipid:lyotrope

In the case of micelles, a core-shell ellipsoid form factor has a fringe after a first minima, which is known to correlate well with overall micelle size [1]. The centre of the peak, $q_c$ can be used for this purpose:

$$d = \frac{2\pi}{q_c} \tag{2}$$

where d is the approximate micelle diameter.

#### S2 Excess lyotrope content

Our method of sample preparation differed from that of Cherezov et al. by using excess lyotropic conditions [2]. To ensure that our results were reproducible and consistent, we measured the excess lyotrope mesophase behaviour using three different lipid:lyotrope weight ratios.

Samples were prepared by weighing a quantity of monoolein, and adding a corresponding weight of lyotrope at a fixed butanediol proportion of 40% v/v as required. The samples were mixed mechanically, transferred to an X-Ray capillary, sealed, and put through 3 freeze-thaw cycles to ensure equilibrium. They were measured for 600s in a q range of 0.015–0.65 $Å^{-1}$.

The azimuthal scattering patterns obtained and plotted in Figure S1 show that beyond an excess point, there is no substantial change in the mesophase behaviour of the monoolein/water/butanediol system between the different weight ratios. The correlation lengths measured for the 0.4, 0.25, and 0.14 %w/w systems were 101 Å, 111 Å, and 101 Å respectively.

### S3 Scattering patterns

## S3.1 MO

Figure S2 shows integrated SAXS patterns for a pure monoolein system with varying proportions of butanediol in the lyotrope.

#### S3.2 Cholesterol

Figures S3 to S6 show integrated SAXS patterns for systems doped with 2.5% mol (Fig. S3), 5% mol (Fig. S4), 7.5% mol (Fig. S5), and 10% mol (Fig. S6) cholesterol.

#### S3.3 DOPE

Figures S7 to S10 show integrated SAXS patterns for systems doped with 2.5% mol (Fig. S7), 5% mol (Fig. S8), 7.5% mol (Fig. S9), and 10% mol (Fig. S10) DOPE.

#### S3.4 DOPC

Figures S11 to S14 show integrated SAXS patterns for systems doped with 2.5% mol (Fig. S11), 5% mol (Fig. S12), 7.5% mol (Fig. S13), and 10% mol (Fig. S14) DOPC.

#### S3.5 DDM

Figures S15 to S18 show integrated SAXS patterns for systems doped with 2.5% mol (Fig. S15), 5% mol (Fig. S16), 7.5% mol (Fig. S17), and 10% mol (Fig. S18) DDM.

#### S3.6 DOPG

Figures S19 to S21 show SAXS patterns measured for systems doped with 1% mol (Fig. S19), 3% mol (Fig. S20), and 5% mol (Fig. S21) DOPG. Some peaks are occaisionally visible, but it was not possible to index them into recognised mesophases in most cases.

##### S3.7 DOPG/cholesterol

Figures S22 to S23 show SAXS patterns for systems doped with 1% mol DOPG and 9% cholesterol (Fig. S22), and 3% mol DOPG and 7% cholesterol (Fig. S23).

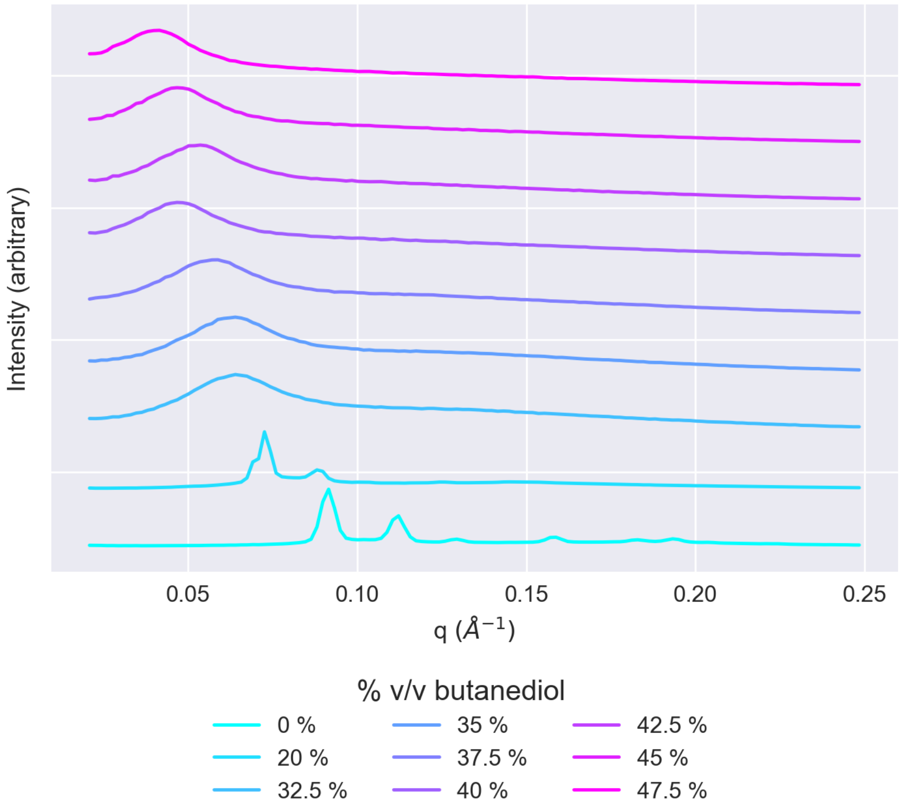


Figure S2: SAXS patterns for a pure MO system hydrated with a lyotrope of butanediol and water. The patterns are ordered with increasing butanediol lyotrope content from bottom to top.

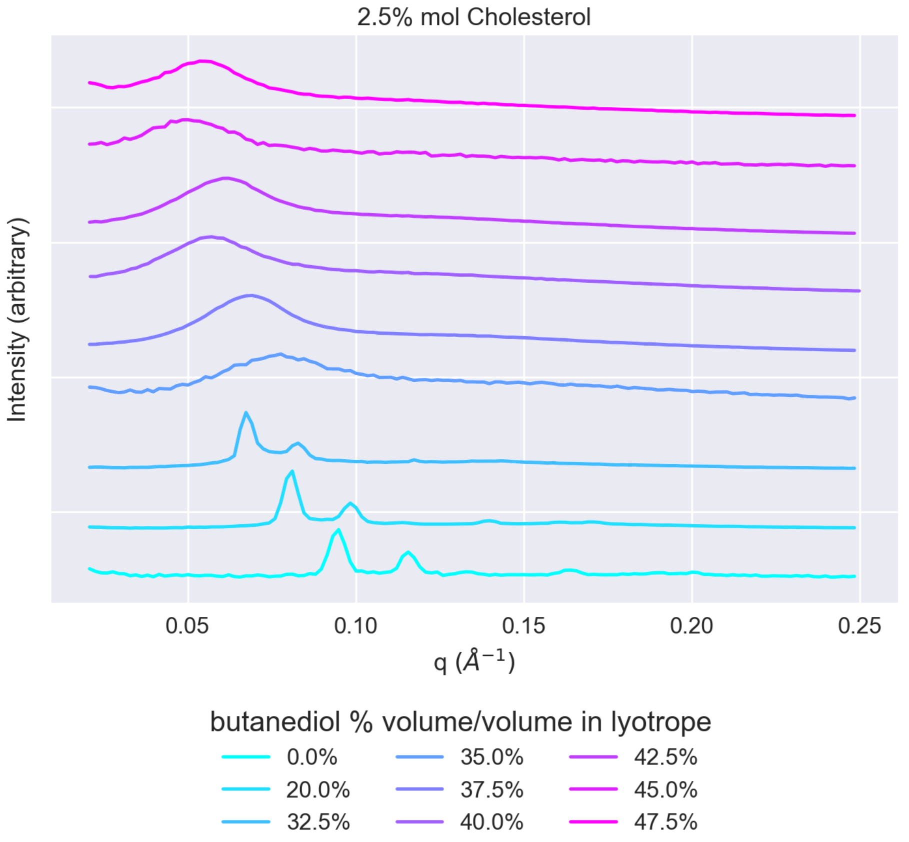


Figure S3: SAXS patterns for systems doped with 2.5% mol cholesterol. The patterns are ordered with increasing butanediol lyotrope content from bottom to top.

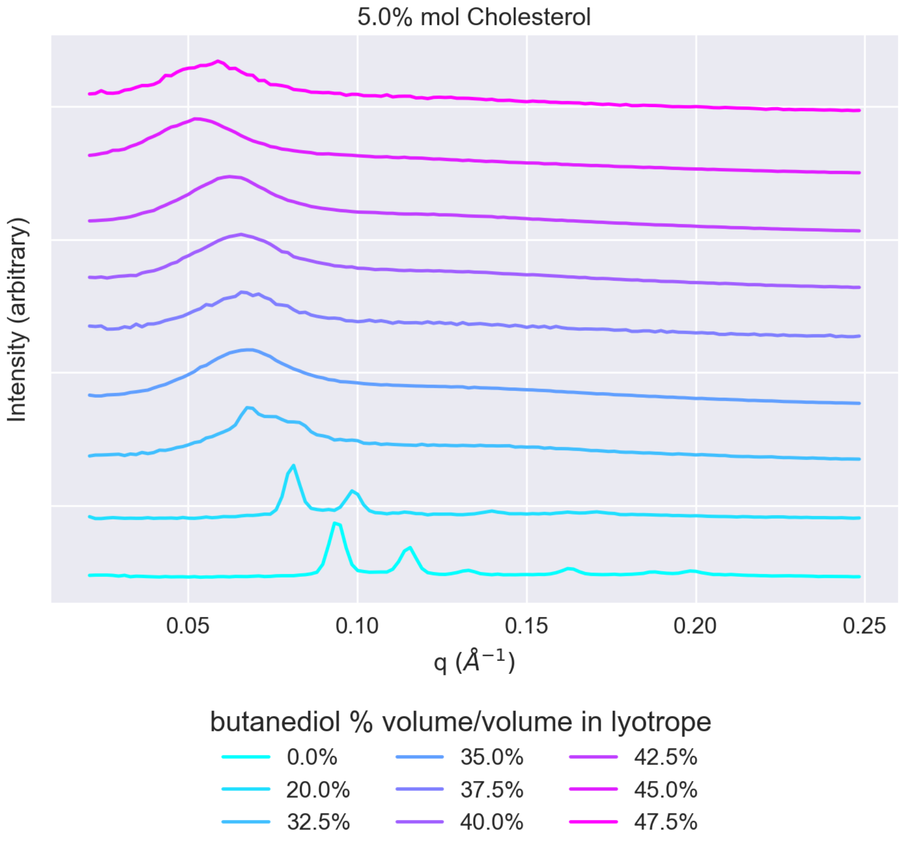


Figure S4: SAXS patterns for systems doped with 5% mol cholesterol. The patterns are ordered with increasing butanediol lyotrope content from bottom to top.

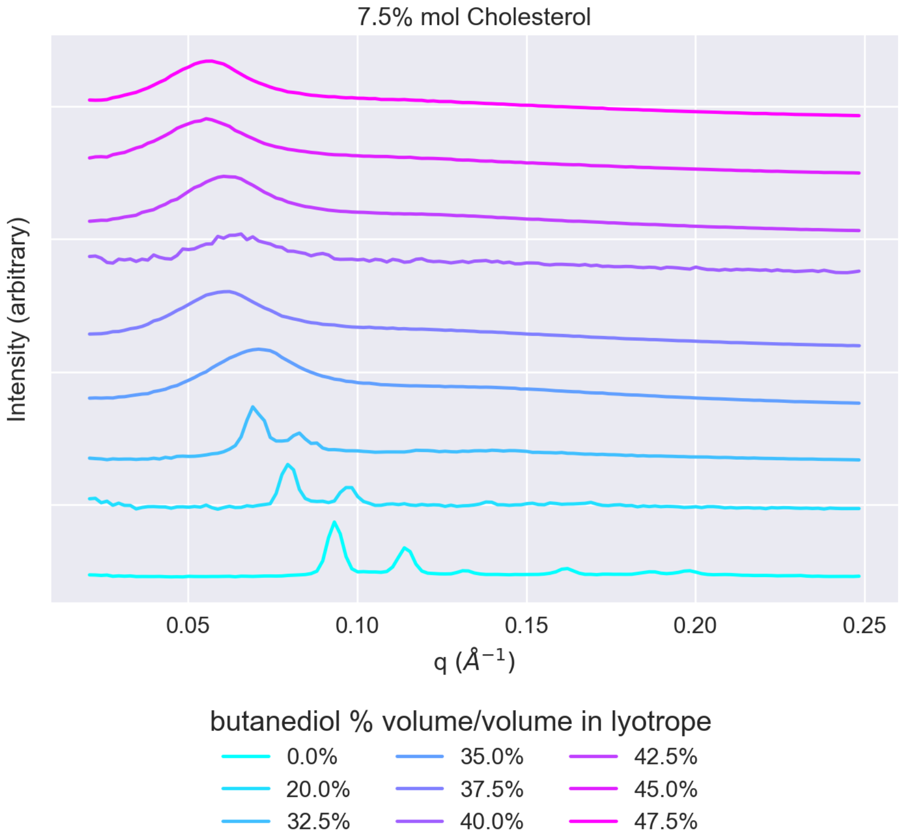


Figure S5: SAXS patterns for systems doped with 7.5% mol cholesterol. The patterns are ordered with increasing butanediol lyotrope content from bottom to top.

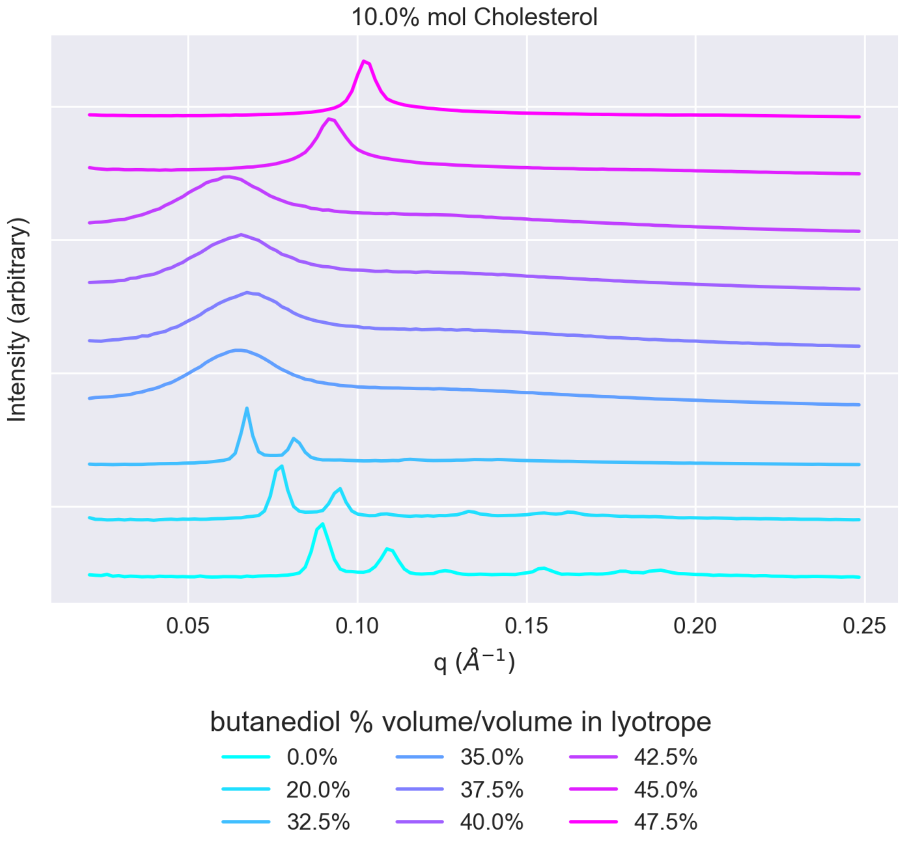


Figure S6: SAXS patterns for systems doped with 10% mol cholesterol. The patterns are ordered with increasing butanediol lyotrope content from bottom to top.

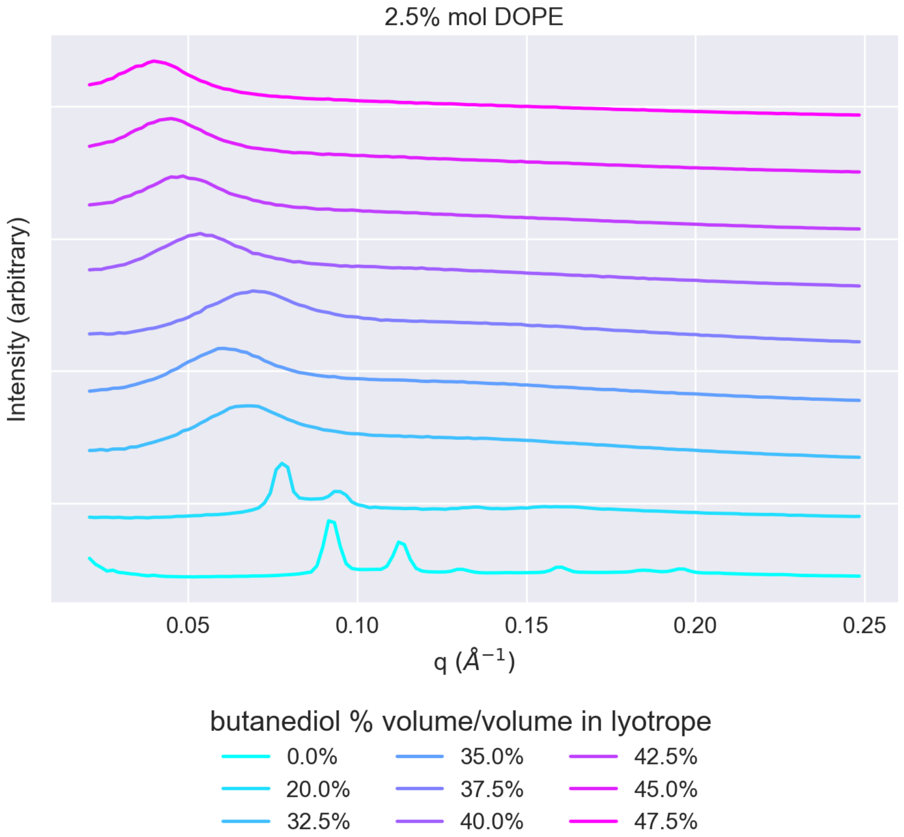


Figure S7: SAXS patterns for systems doped with 2.5% mol DOPE. The patterns are ordered with increasing butanediol lyotrope content from bottom to top.

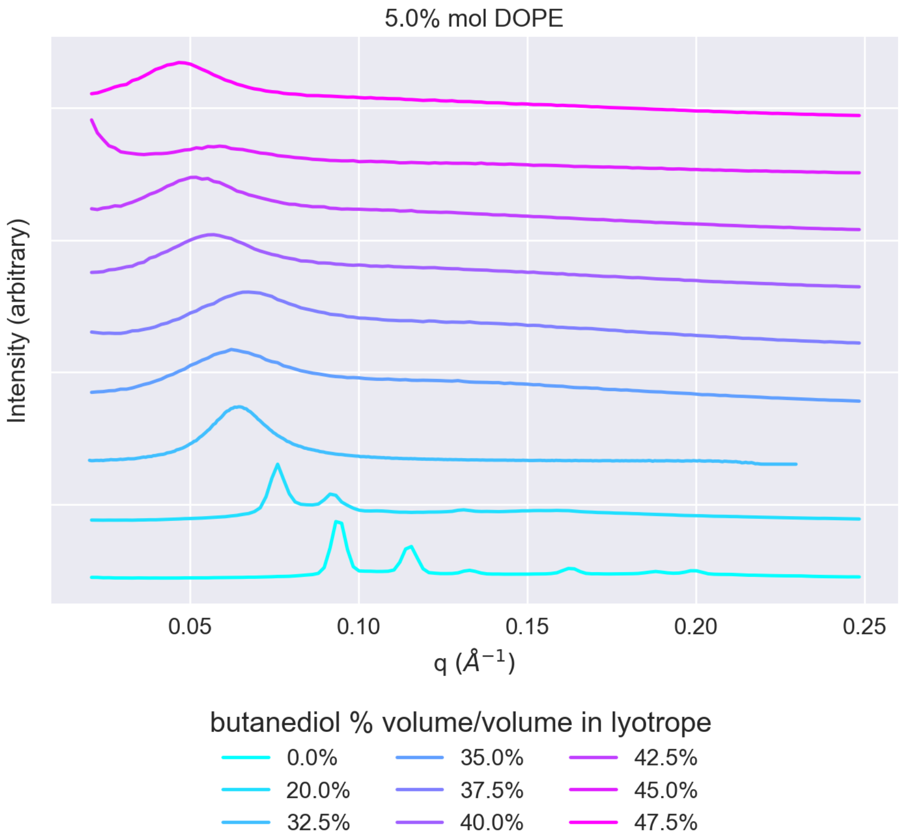


Figure S8: SAXS patterns for systems doped with 5% mol DOPE. The patterns are ordered with increasing butanediol lyotrope content from bottom to top.

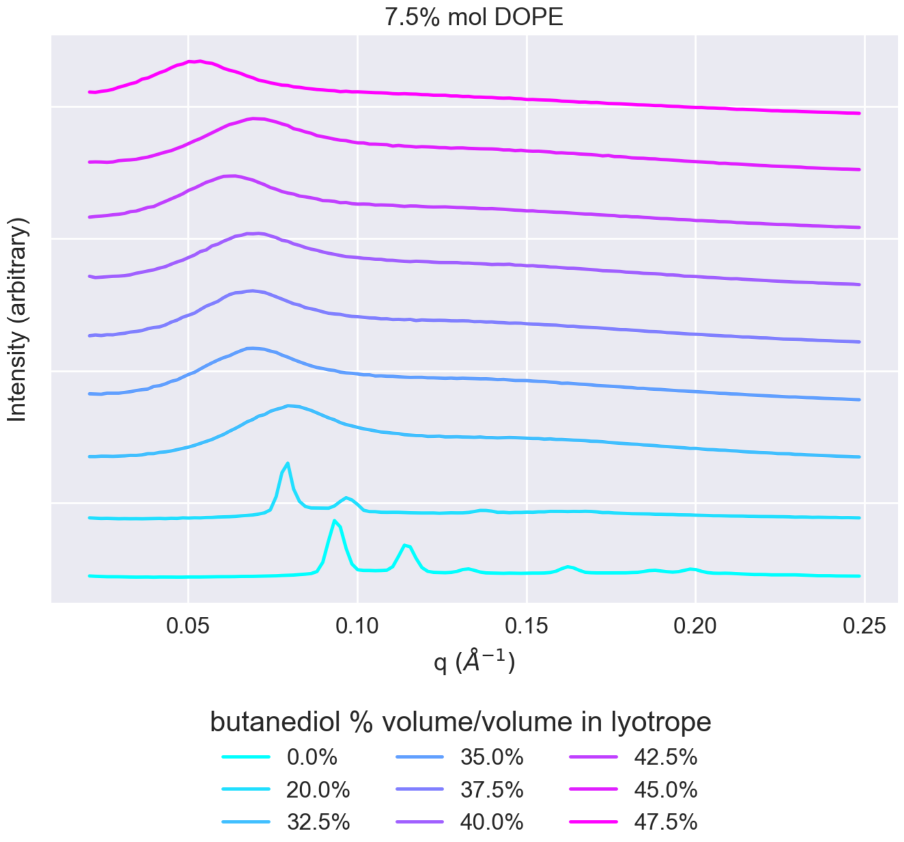


Figure S9: SAXS patterns for systems doped with 7.5% mol DOPE. The patterns are ordered with increasing butanediol lyotrope content from bottom to top.

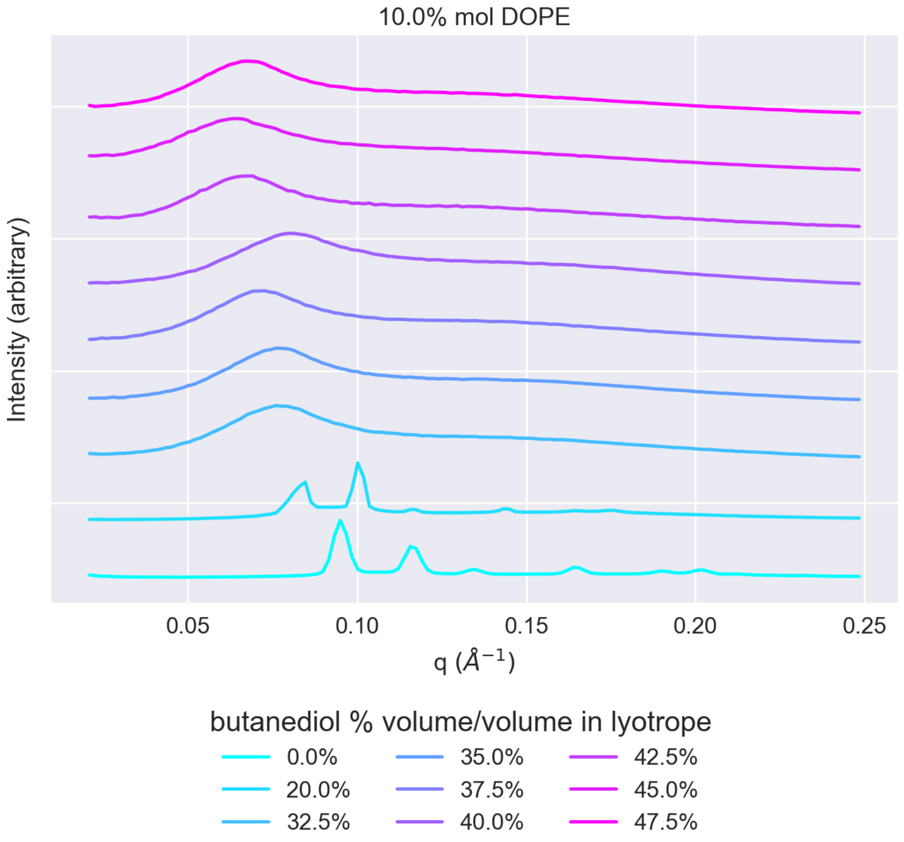


Figure S10: SAXS patterns for systems doped with 10% mol DOPE. The patterns are ordered with increasing butanediol lyotrope content from bottom to top.

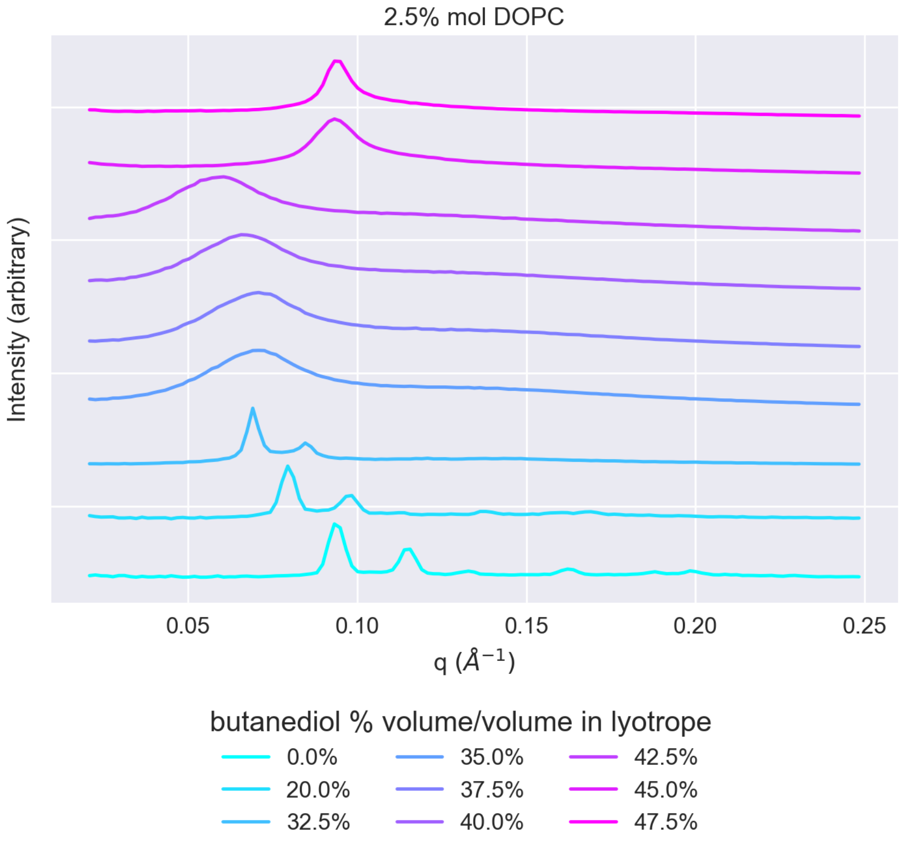


Figure S11: SAXS patterns for systems doped with 2.5% mol DOPC. The patterns are ordered with increasing butanediol lyotrope content from bottom to top.

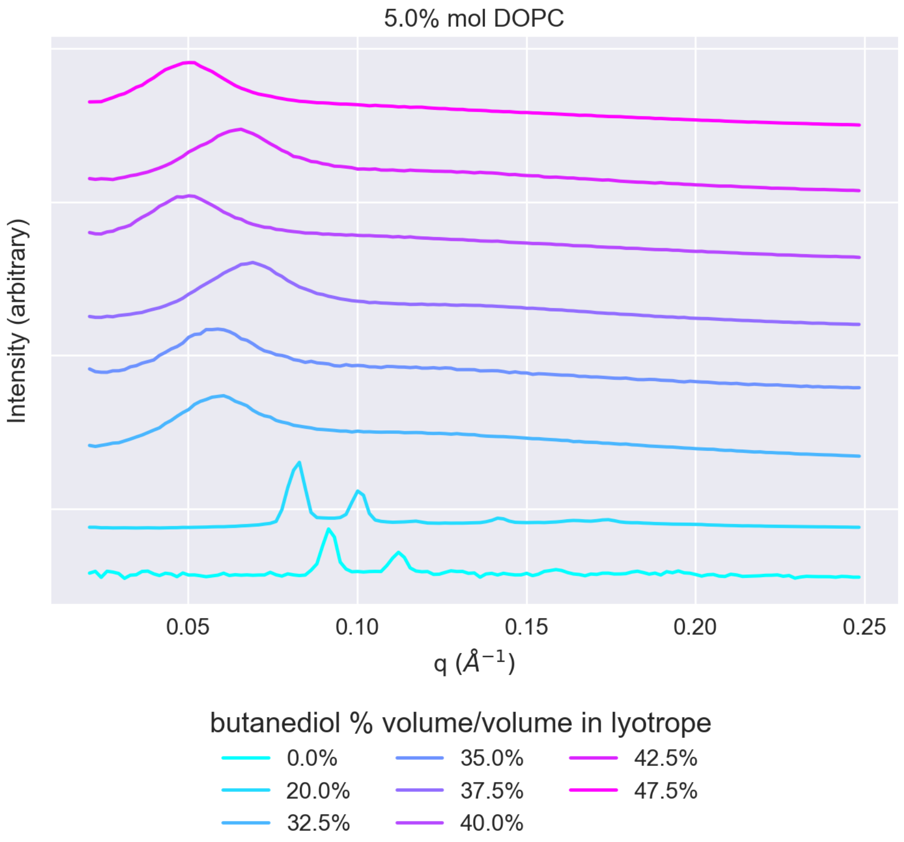


Figure S12: SAXS patterns for systems doped with 5% mol DOPC. The patterns are ordered with increasing butanediol lyotrope content from bottom to top.

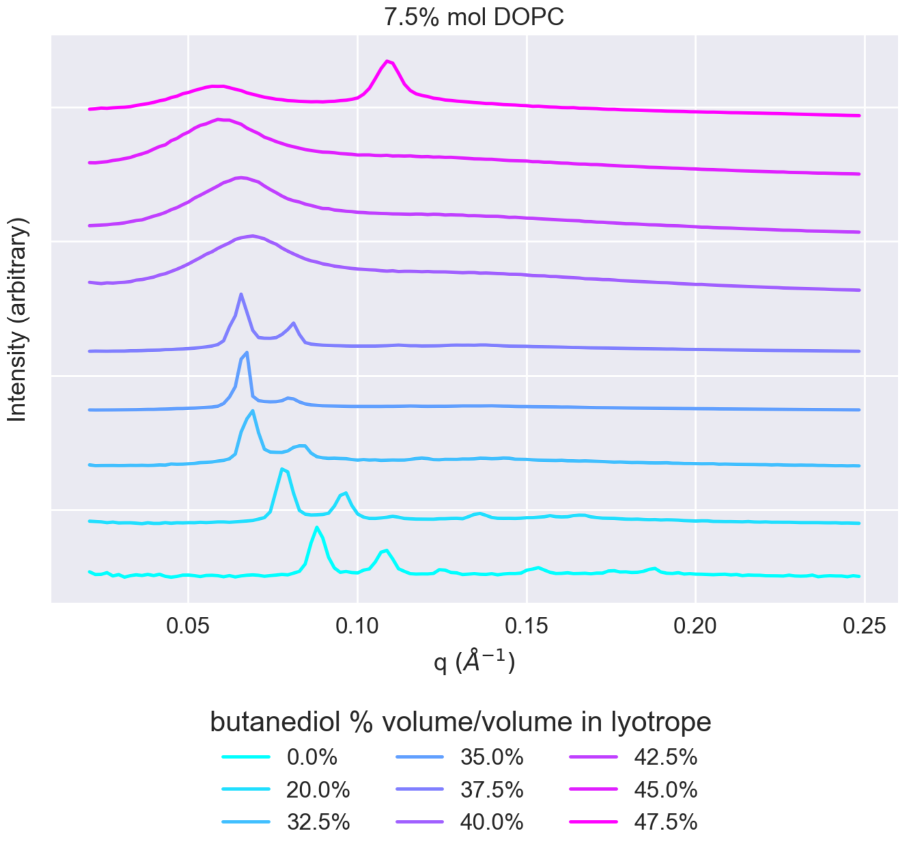


Figure S13: SAXS patterns for systems doped with 7.5% mol DOPC. The patterns are ordered with increasing butanediol lyotrope content from bottom to top.

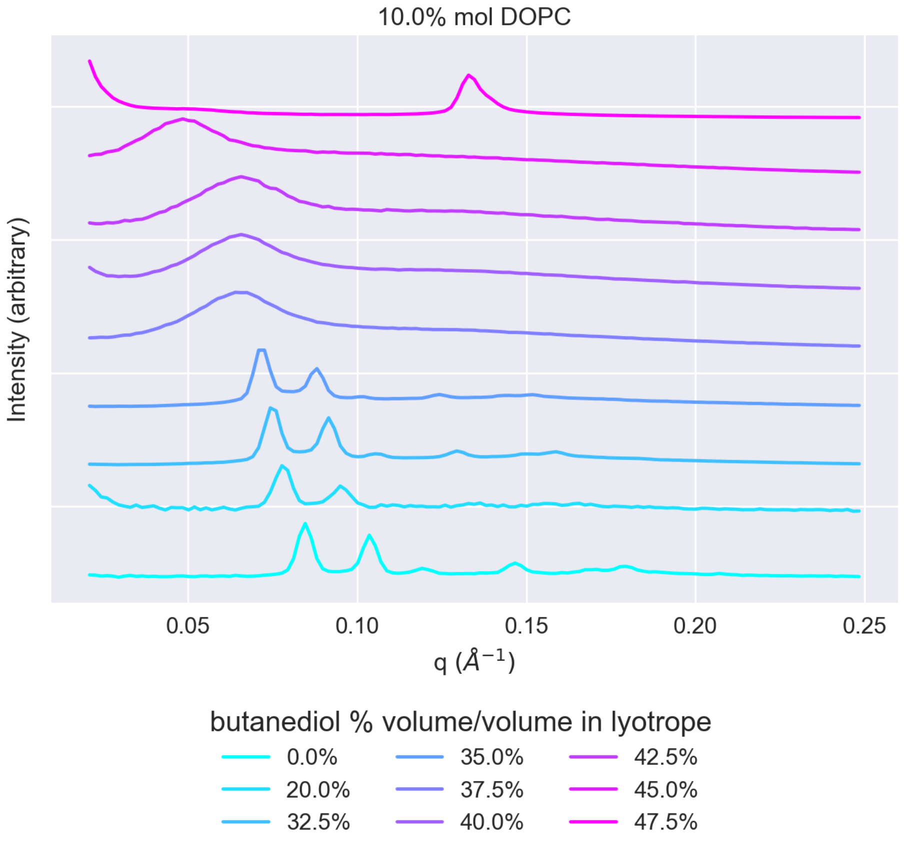


Figure S14: SAXS patterns for systems doped with 10% mol DOPC. The patterns are ordered with increasing butanediol lyotrope content from bottom to top.

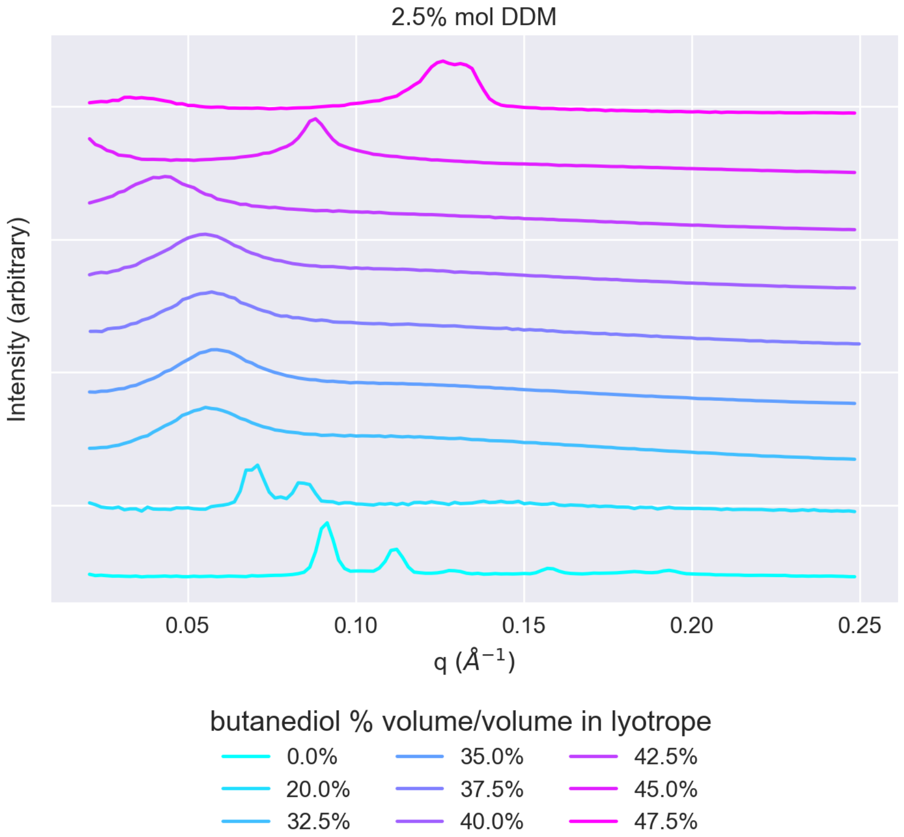


Figure S15: SAXS patterns for systems doped with 2.5% mol DDM. The patterns are ordered with increasing butanediol lyotrope content from bottom to top.

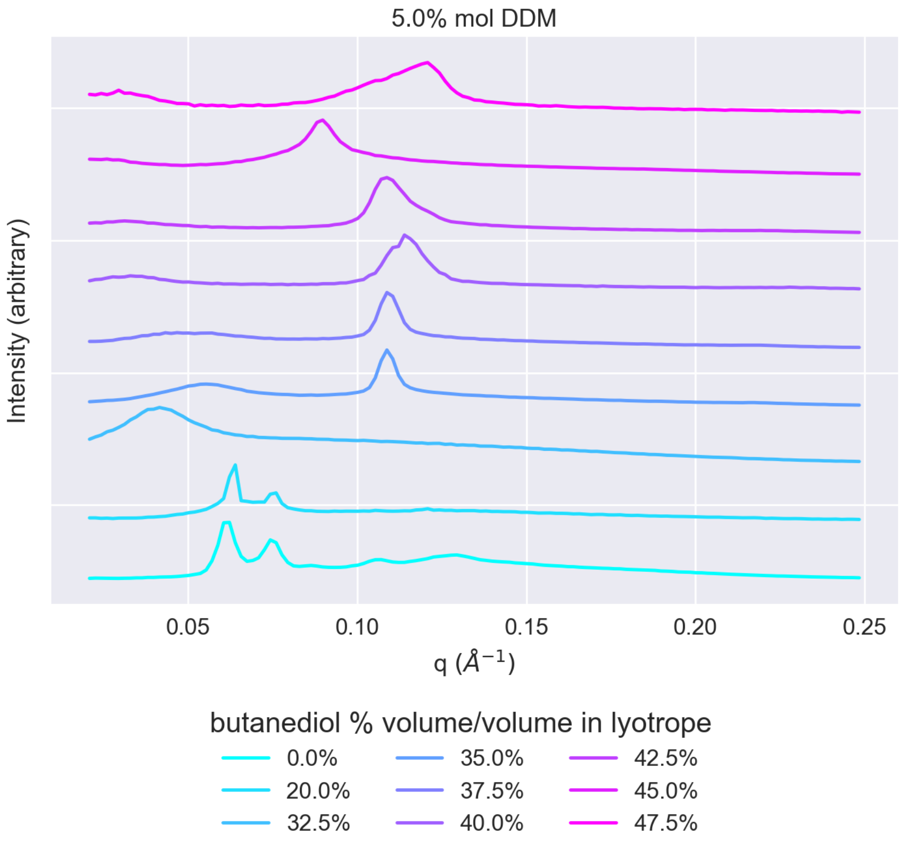


Figure S16: SAXS patterns for systems doped with 5% mol DDM. The patterns are ordered with increasing butanediol lyotrope content from bottom to top.

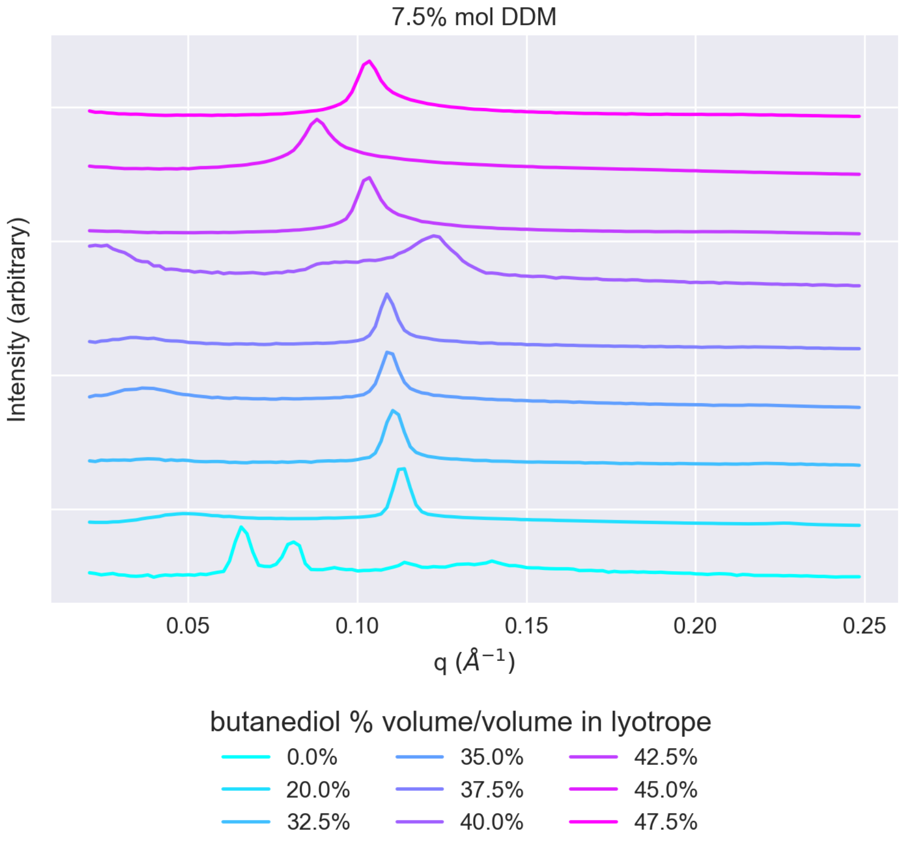


Figure S17: SAXS patterns for systems doped with 7.5% mol DDM. The patterns are ordered with increasing butanediol lyotrope content from bottom to top.

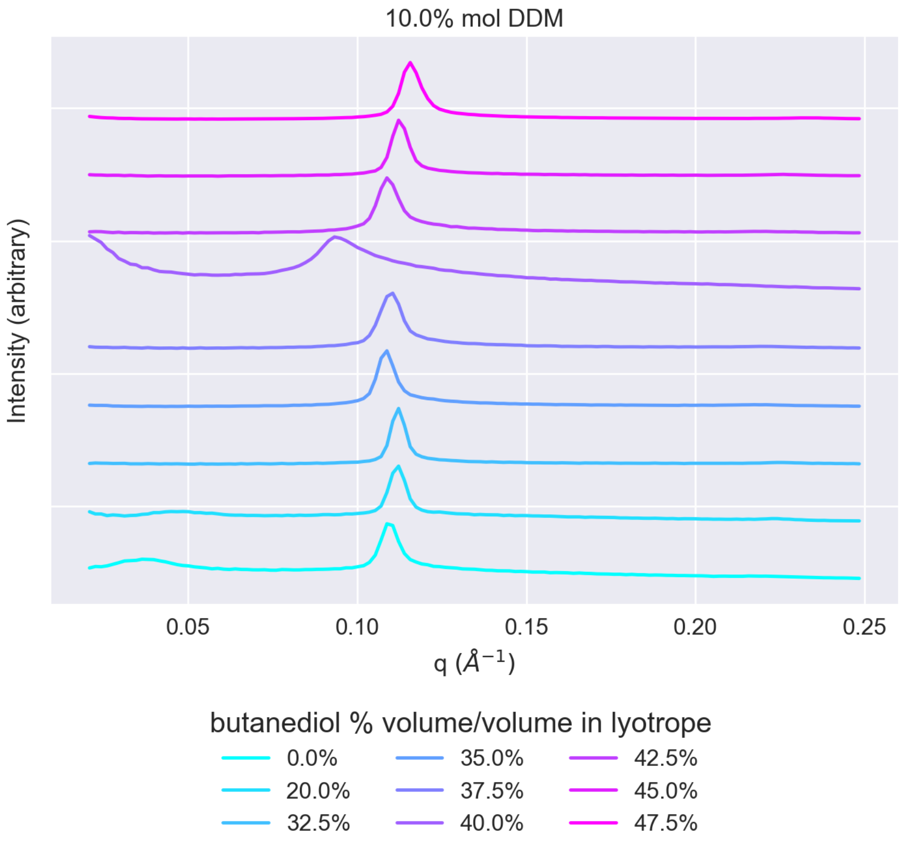


Figure S18: SAXS patterns for systems doped with 10% mol DDM. The patterns are ordered with increasing butanediol lyotrope content from bottom to top.

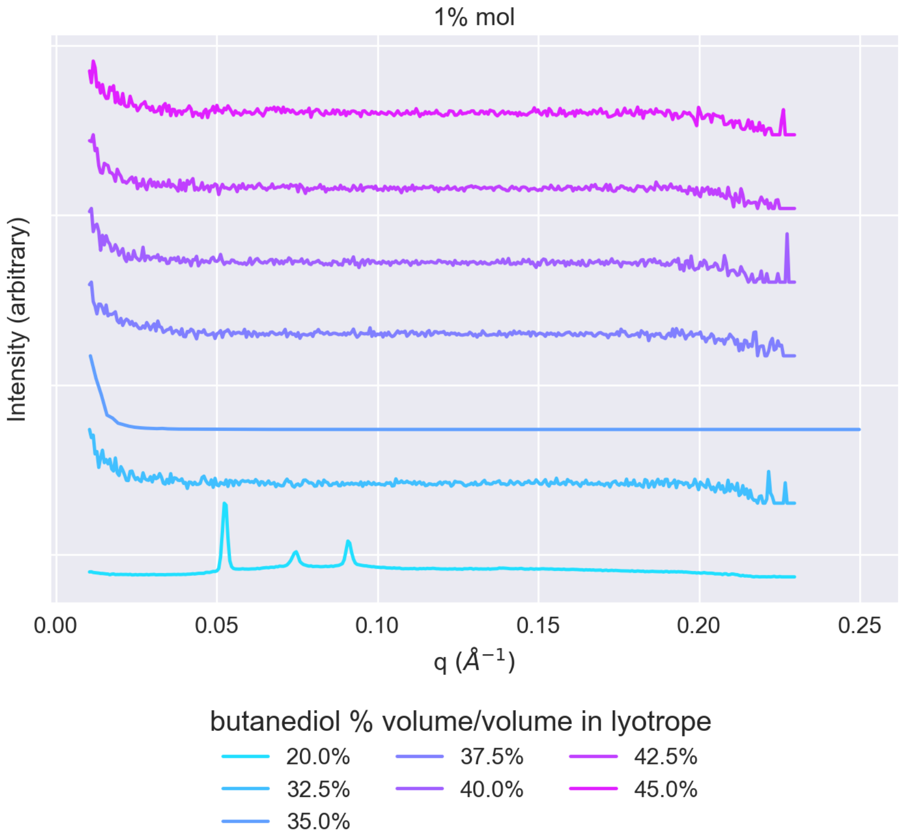


Figure S19: SAXS patterns for systems doped with 1% mol DOPG.

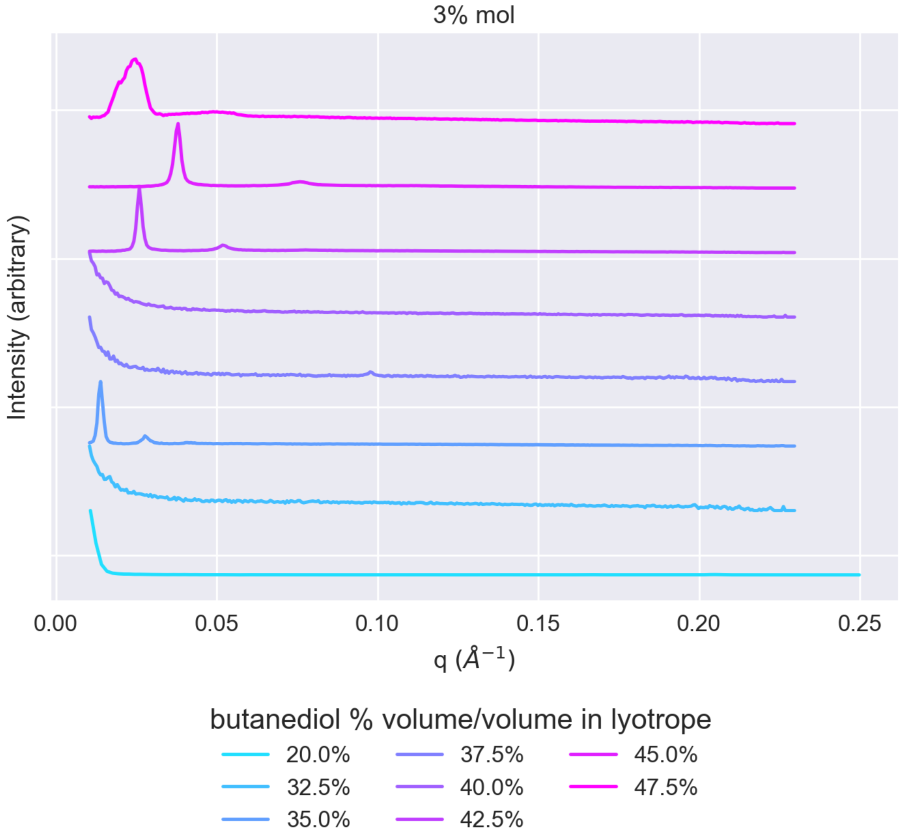


Figure S20: SAXS patterns for systems doped with 3% mol DOPG.

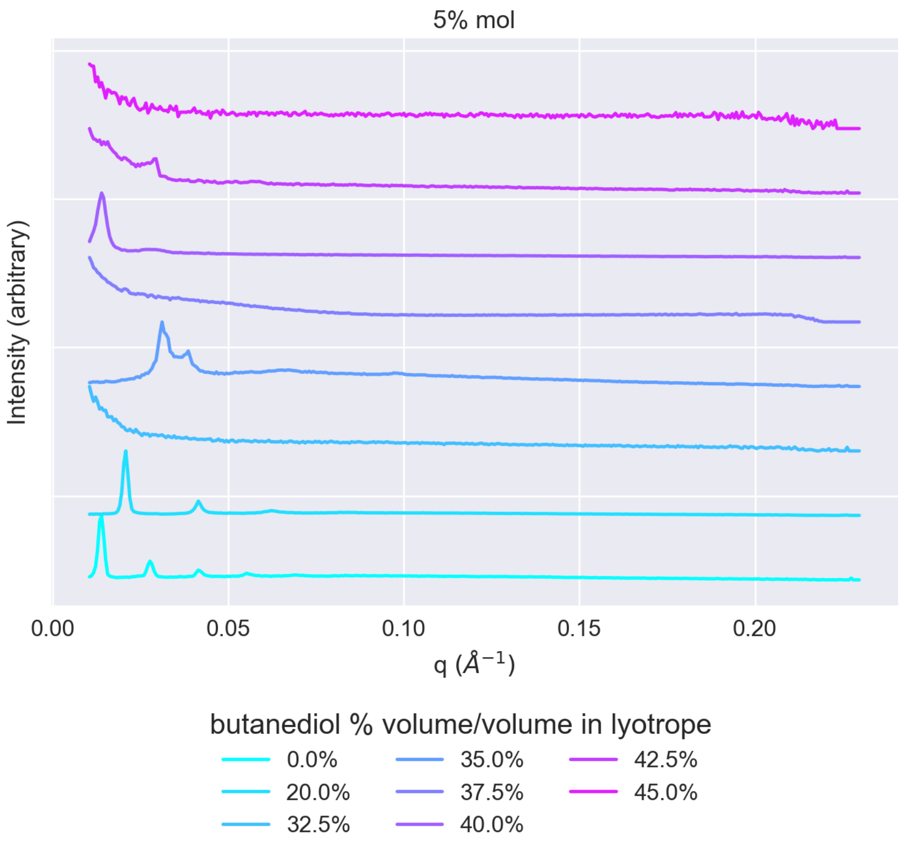


Figure S21: SAXS patterns for systems doped with 5% mol DOPG.

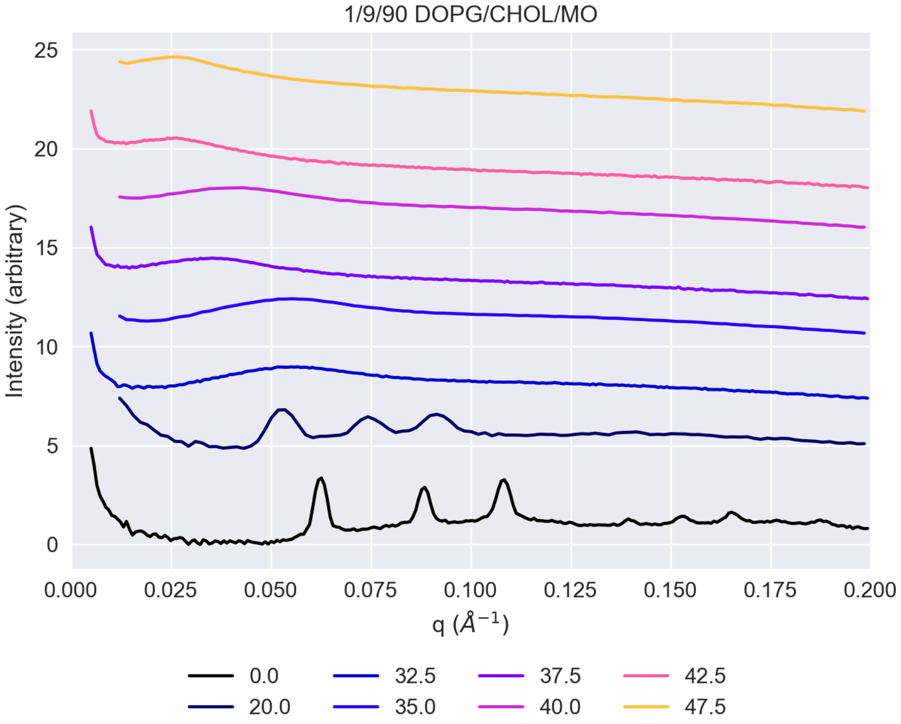


Figure S22: SAXS patterns for systems doped with 1% mol DOPG, 9% mol cholesterol, and 90% mol MO.

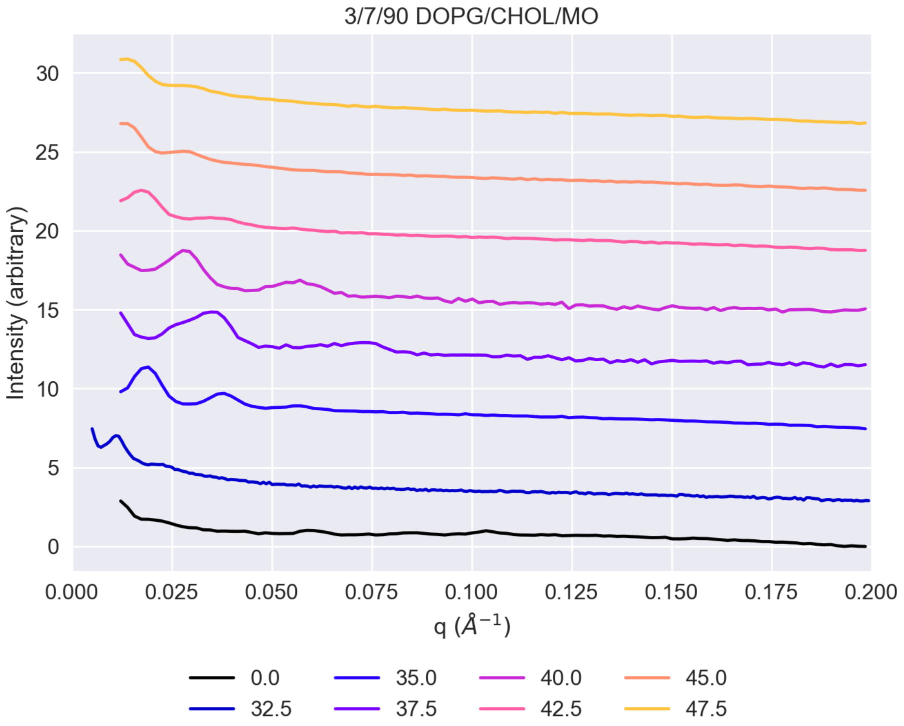


Figure S23: SAXS patterns for systems doped with 3% mol DOPG, 7% mol cholesterol, and 90% mol MO.
